# Determinants of Water Source Use, Quality of Water, Sanitation and Hygiene Perceptions among Urban Households in North-West Ethiopia: A Cross-Sectional Study

**DOI:** 10.1101/2020.09.09.289199

**Authors:** Shewayiref Geremew Gebremichael, Emebet Yismaw, Belete Dejen, Adeladilew Dires

**Affiliations:** Department of Statistics, Debre Tabor University, Debre Tabor, Amhara, Ethiopia; Department of Mathematics, Debre Tabor University, Debre Tabor, Amhara, Ethiopia

**Keywords:** Water Source, Households, Perception, Quality, Sanitation

## Abstract

**Background:** Clean water is an essential element for human health, wellbeing, and prosperity. Every human being has the right to access safe drinking water. But, in now day, due to rapid population growth, illiteracy, lack of sustainable development, and climate change; it still faces a global challenge for about one billion people in the developing nation. The discontinuity of drinking water supply puts in force households either to use unsafe water storage materials or to use water from unimproved sources. This study aimed to identify the determinants of water source types, use, quality of water, and sanitation perception of physical parameters among urban households in North-West Ethiopia.

**Methods:** A community-based cross-sectional study was conducted among households from February to March 2019. An interview-based pre-tested and structured questionnaire was used to collect the data. Data collection samples were selected randomly and proportional to each kebeles’ households. MS Excel and R Version 3.6.2 was used to enter and analyze the data; respectively. Descriptive statistics using frequencies and percentages were used to explain the sample data concerning the predictor variable. Both bivariate and multivariate logistic regressions were used to assess the association between the independent and the response variables.

**Results:** Four hundred eighteen (418) households have participated. Based on the study undertaken, 78.95% of households used improved and 21.05% of households used unimproved drinking water sources. Households drinking water sources are significantly associated with age of participant (x^2^ = 20.392, df=3), educational status (x^2^ = 19.358, df=4), source of income (x^2^ = 21.777, df=3), monthly income (x^2^ = 13.322, df=3), availability of additional facilities (x^2^ = 98.144, df=7), cleanness status (x^2^ =42.979, df=4), scarcity of water (x^2^ = 5.1388, df=1) and family size (x^2^ = 9.934, df=2). The logistic regression analysis also indicated as those factors are significantly determined (p 0.05) the water source types used by households. Factors such as availability of toilet facility, household member type, and sex of head of the household are not significantly associated with the drinking water sources.

**Conclusion:** The study showed that being an older age group of the head of the household, being government employer, merchant and self-employed, being a higher income group, the presence of all facilities in the area, lived in a clean surrounding and lower family size are the determinant factors of using drinking water from improved sources. Therefore; the local, regional, and national governments and other supporting organizations shall improve the accessibility and adequacy of drinking water from improved sources through short and long time plans for the well-being of the community in the area.

## 1 BACKGROUND

Clean water is an essential element for human health, wellbeing, and prosperity [1]. Every human being has the right to access safe drinking water. But, in now day, due to rapid population growth, illiteracy, lack of sustainable development, and climate change (drought and poverty) are still face a global challenge. Currently, about one billion people, who live in the developing world, don’t have access to safe and adequate drinking water [2].

Water found from either improved or unimproved water sources. An improved water source is a term used to categorize certain types or levels of water supply for monitoring purposes. It is defined as a type of water source that, by nature of its construction or through active intervention, is likely to be protected from outside contamination, in particular from contamination with fecal matter [3]. Improved water sources are those that have the potential to deliver safe water by nature of their design and construction [3–4]. These include piped supplies (such as households with tap water in their dwelling, yard or plot; or public stand posts) and non-piped supplies (such as boreholes, protected wells and springs, rainwater, and packaged or delivered water). Between 2000 and 2015, the population using piped supplies increased from 3.5 billion to 4.7 billion, while the population using non-piped supplies increased from 1.7 billion to 2.1 billion. Globally, two out of five people in rural areas and four out of five people in urban areas use piped supplies [4]. The opposite of “improved water source” has been termed “unimproved water source”, based on the JMP [5] definitions.

A report of Ethiopia Socioeconomic Survey-2016 [6] also classifies the water sources as improved and unimproved. Improved water sources are included piped water from any location, tube wells or boreholes, protected dug wells, protected springs, rainwater, bottled water, and water delivered by tanker truck or cart. Similarly, unimproved water sources are water collected from unprotected dug wells, unprotected springs, and surface water.

About 748 million people{mostly the poor and marginalized-there is a scarcity of using an improved water source supply and of these, almost a quarter (173 million) rely on untreated surface water, and over 90% live in rural areas [7]. About 547 million people didn’t have an improved drinking water supply in 2015.

A study conducted by Water.org [8], found that 42% of the population has access to a clean water supply and only 11% of that number has access to adequate sanitation services.

In Africa, only 60% of the population has access to improved sanitation services, but the situation is worse in rural areas, in which below half (45%) of the rural population has access to improved sanitation services. According to WHO, 2011 report, individuals with no access to improved sanitation are forced to defecate in open fields, in rivers, or near areas where children play and food is prepared [9].

The Ethiopian Demographic and Health Survey (EDHS) (CSA and ICF, 2016) [10] reported that 97% of urban households in Ethiopia have access to an improved source of drinking water and in rural areas, only 57% improved water access. Nevertheless, no reliable information is available on the readability of drinking water quality reports for further illustration [11].

Based on reports, Ethiopia is the country with the worst of all water quality problems in the world. It has the lowest water supply (42%) and sanitation coverage (28%) in sub-Saharan countries [12]. Ethiopia is considered one of the poorest sanitation and drinking water infrastructure [13]. Beyene et al., (2015) [14] report shows that about 52.1% of the population has been using unimproved sanitation facilities while 36% of them practiced open defecation. In Ethiopia due to the discontinuity of drinking water supply, it affects the distribution of water to the community in need [15].

According to the data from the Joint Monitoring Programme for Water Supply and Sanitation of WHO/UNICEF, which are in turn based on data from various national surveys including the 2005 Ethiopian Demographic and Health Survey (EDHS), access to an improved water source and improved sanitation was estimated as 38% for improved water supply (98% for urban areas and 26% for rural areas), and 12% for improved sanitation (29% in urban areas, 8% in rural areas) [5].

The low quality of water and lack of accessibility of it have been faced different challenges in the household members. Such as: due to the lack of access to water in many rural children, especially females are put to work collecting water each morning and to help their families [16,17]; the burden of water collection does not fall equally on all household members [6]. Thus; 75 percent are female and 25 percent are male. The report [6] also added the gender breakdown is consistent for both urban and rural areas. The responsibility of collecting water-primarily falls on women, sons, or daughters of the household. Globally; women (64%), men (24%), girls (8%), and boys (4%) share the burden of collecting water [18]. Due to the presence of a burden on children, only 45% of kids attend primary education in Ethiopia [16], and also, water and sanitation-related sicknesses put severe burdens on health services and keep children out of school [19]. Younger household members are more likely to collect water, but this differs by place of residence; while only 22 percent of those who collect water in urban areas are children (ages 7 to 14); in rural areas, nearly 37 percent of water collectors are children [20].

The discontinuity of drinking water supply obligate households to use water storage material or to use water from unimproved sources. In the study area, the irregularity of water supply is observed, due to this the community use water storage materials. Thus, water stored in unprotected materials (such as unsafe Pot, Rotto, Jerikan, other plastic materials) for a long time might be contaminated and cause water-borne diseases. The water from unimproved water sources might be contaminated with animals, floods, and specks of dust through wind and human wastes. This may perhaps cause illness to humans.

The objective of this study is to assess the water sources types households’ use for fetching water; identify the associated factors of using improved sources of water, and to determine the quality of water and sanitation perception of physical parameters in urban households in North-West Ethiopia. This article provides data on access, usage, and practices of water sources among urban households in Northwest Ethiopia. This article provides a valuable insight into access to safe water and consequent socioeconomic conditions such as income, distances, and family size that can become a barrier to access.

To remind the reader that the data used in this study is based on perceptions of households and no verifiable water quality measures were undertaken by the investigators.

## 2 METHODS

### 2.1 Study Design, Area and Period

Community-based cross-sectional study design was conducted from February to March 2019. Debre Tabor Town is the Zonal Administration center of South Gondar Zone, located 666 km North-West from the capital city of Ethiopia, Addis Ababa. Debre Tabor is found 103 km East away from Bahir Dar, the regional town of Amhara National Regional State. It is located about 100 km South-East of Gondar and 50 km East of Lake Tana. Geographically, Debre Tabor has a latitude and longitude of 11^0^51^0^N and 38^0^1^0^E; respectively and an elevation of 2,706 meters above sea level. The average temperature of the town is 14.8^0^C and the average annual rainfall is 1497 mm. The town is known for its cold weather condition due to the presence of Mount Guna nearby. Based on the town official report, the population of the town is expected to be 87,627 (2019 projected population); of this number 49,535 household members are users of tap water in 2019, but the left 38,092 household members are not tapped, water users. There are agricultural farms in and around the city which are irrigated with river water and dug water. Animal farming in the town is very common using dug water sources. Drinking water of the town comes from large reservoirs located in its surroundings, Farta woreda, which is one of the administrative woredas in South Gondar Zonal Administration. (Figure 1)

**Figure 1:**
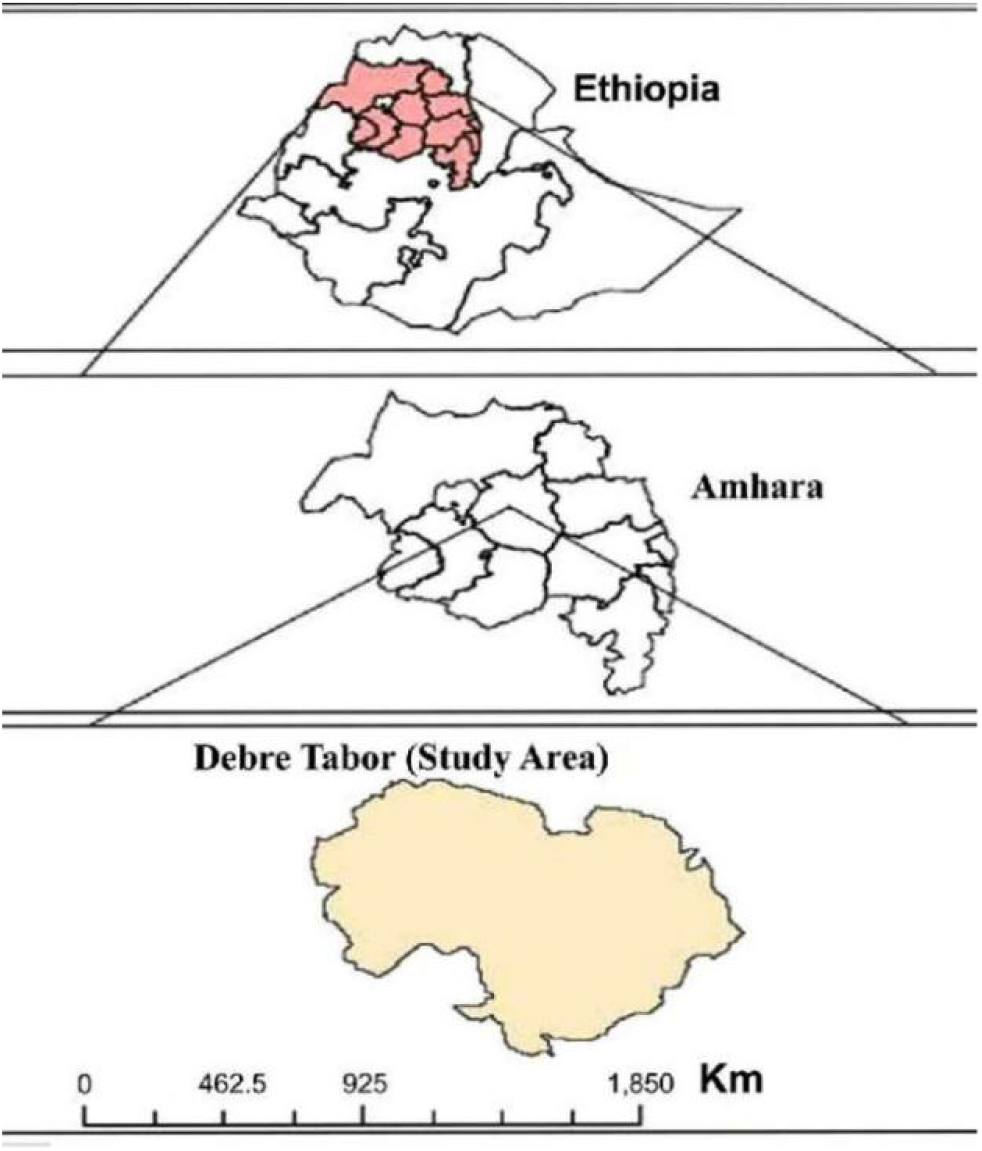
Location map of the study area

There are a lot of wells, which were extracted to supplement the domestic water requirement in Debre Tabor Town. Based on the previous evidence the households of Debre Tabor town get pipe water supply only once in a week [21].

### 2.2 Sample Size Determination

The town has about 17,526 households and of which study population is made up of families that reside in the town, which comprised men, women, and children, all of which are in different age groups. The average family size of the town was computed about 4.53 per household (After a pilot survey was conducted in January 2019).

For the household survey, samples were decided to select by using the sample size determination equation of Cochran (1977) [22].

The study used a single proportion formula, 95% confidence interval, the marginal error of 5%, and the non-response rate of 10%;

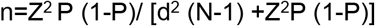

Where;

Z = 95% confidence limit (z-value at α=0.05 is 1.96)
N = Number of households in Debre Tabor town =17526
P=0.5, 1-P=0.5, D=Marginal error or degree of accuracy =0.05 n=380+38=418
Total sample size=418. However, a sample size of 418 was used to eliminate any errors.

The town has 6 kebeles. The samples were selected randomly and taken proportionally from each kebeles and sub-kebeles.

### 2.3 Sampling Method and Sampling Procedure

All households who were lived in the study area at the time of data collection and fulfill the inclusion criteria were included as the study population. Data collection sites were demarcated into 6 kebeles. The samples were selected randomly and taken proportionally from each kebeles and sub-kebeles.

### 2.4 Data Collection Tools and Techniques

The primary source of data was employed. The primary data gathering was including, household survey questionnaires (on paper) and personal observation. The content of the questionnaire was checked by public health professionals, who have had a profound experience in the area. The method of data collection was by investigator administered questionnaire. The investigators/researchers (people who speak the Amharic language) administered a questionnaire that was administered to participants regardless of their educational level. Seven data collectors were selected for data collection and two supervisors were assigned.

The questionnaire consisted of five sections namely; section I: socio-demographic data, section II: Sources of income, section III: water source observation, section IV: household water use and section V: Water quality and sanitation perception. There were key informant interviews and verification of the facilities using a checklist.

A detailed questionnaire was prepared in the native language of the households (Amharic) and included over 50 questions. A multiple-choice format was used to answer the majority of the questions. House-hold characteristics, such as the number of family size, educational level, monthly income of the household, type of occupation, sources of water, sanitation and hygiene, and awareness about household drinking water were included.

### 2.5 Study variables

The variables included in this study are taken based on perceptions of households and not verifiable water quality measures were undertaken.

#### 2.5.1 Dependent variables

The dependent variable is sources of drinking water (improved, unimproved). Based on WHO guidelines: ‘improved water sources’ consisted of piped water (into dwelling, yard/plot, and public tap/stand-pipe), tube-well/ bore-hole, protected well, protected spring, and rainwater. Bottled water was included as an improved water source if the household used another improved water source for other purposes, such as hand-washing and cooking. Unprotected wells, unprotected springs, tanker trucks, a cart with tank/drum, and surface water was considered as ‘unimproved water sources’.

#### 2.5.2 Independent variables

The independent variables included in this study are demographic, socio-economic, and sanitation and hygiene perception characteristics.

✓ The perceptions of households using water from unimproved sources(income, distance from home to water source, the presence of alternative water source, quality of water perception, adequacy of water, waiting time to fetch water, personal interest, and other reasons).
✓ The presence of scarcity of water in the area (yes, no).
✓ The reason that households believe the presence of scarcity of water has occurred (government weakness, a local people problem, and both local people and government problems).
✓ The perception of households of the water they consume safety status (not safe at all, somewhat safe, partially safe, safe, and highly safe).
✓ Households’ perception of the indicator of water quality (color, taste, odor, disease attack, and the presence of all the cases).
✓ Households’ perception of the taste, odor, and color of the water from the improved and unimproved sources were the same as (yes, no).
✓ The causes of water quality problem households perceive (water-containing material, animal wastes, human wastes, flood, and all cases).
✓ Treatment measures households had undertaken during unsafe drinking water (no use at all, boiling, sedimentation, using wuha agar, other methods, and use all measures).
✓ The number of times household members had got sickness due to water related disease and visited health centers for physician assistance with in one year before the survey time (not at all, once, twice, three times, more than three times).
✓ The presence of health extension workers assistance (yes, no) and the number of times the family was visited by health extension workers with one year before the survey (not at all, once, twice, three times, more than three times).
✓ Previous Participation of household members in educational and awareness activities about sanitation and hygiene in their locality (yes, no). The presence of latrine facility in the household’s compound (yes, no) and who were used the latrine (wife, husband, children, and all families, except children). The place household members were defecate (public, neighbor, open el, own toilet), and the presence of the culture of households washing hand after defecation (yes, no).

### 2.6 Data Quality Assurance

A pilot survey (pre-test) was incorporated to recheck the questionnaire and for sample size determination. Before the main survey, mini-survey (pilot survey) was done on 30 households outside of the study area (Bahir Dar city) to avoid exclusion of households who were in the study area due to pre-test. After pre-test questionnaire was done; question order, alternative option, skip pattern and overlapping option were amended. Supervisors and data collectors were trained by principal investigator. During data collection, data collectors were supervised by supervisors in close up.

### 2.7 Operational Definitions

➢ Awareness: Understanding the implication and becoming conscious of conditions and practices concerning many things including hygiene and health.
➢ Improved sanitation: A sanitation system that is connected to public sewer, septic tank and a pours toilet/latrine. For a simple pit latrine, it implies the use of slab and ventilated improved latrine.
➢ Safe water: A water system that is well protected from contamination sources, treated with chemicals and used in ways that prevent contamination.
➢ Sanitation: Act of cleanliness and containment of waste products to make the living and working environment free from matters that affect health and wellbeing.

### 2.8 Data Processing and Analysis

The collected data were coded and entered into MS excel, cleaned, stored and exported into R version 3.6.2 for analysis. Any error occurred during data entry was corrected by revising the original completed questionnaire. Descriptive statistics using frequencies and percentages were used to explain the sample data concerning the predictor variable. Both bivariable and multivariable logistic regression estimates were used to assess the association between the independent and the response variables. During the bivariate analysis p-value 0.2 were included to the multivariate analysis to control the association of confounding variables with that of the response variable. In the multivariable analysis of binary logistic regression, variables with a p-value of 0.05 and 95% confidence interval were considered as statistically significant.

## 3 RESULTS

The summary statistics for different explanatory variables (Table 1) and the cross-tabulations among the types of water sources used by the households and the demographic, economic, sanitation and hygiene perception of households (Table 2) are presented. The results of chi-square tests of association and logistic regression are presented under (Table 2).

**Table 1:**
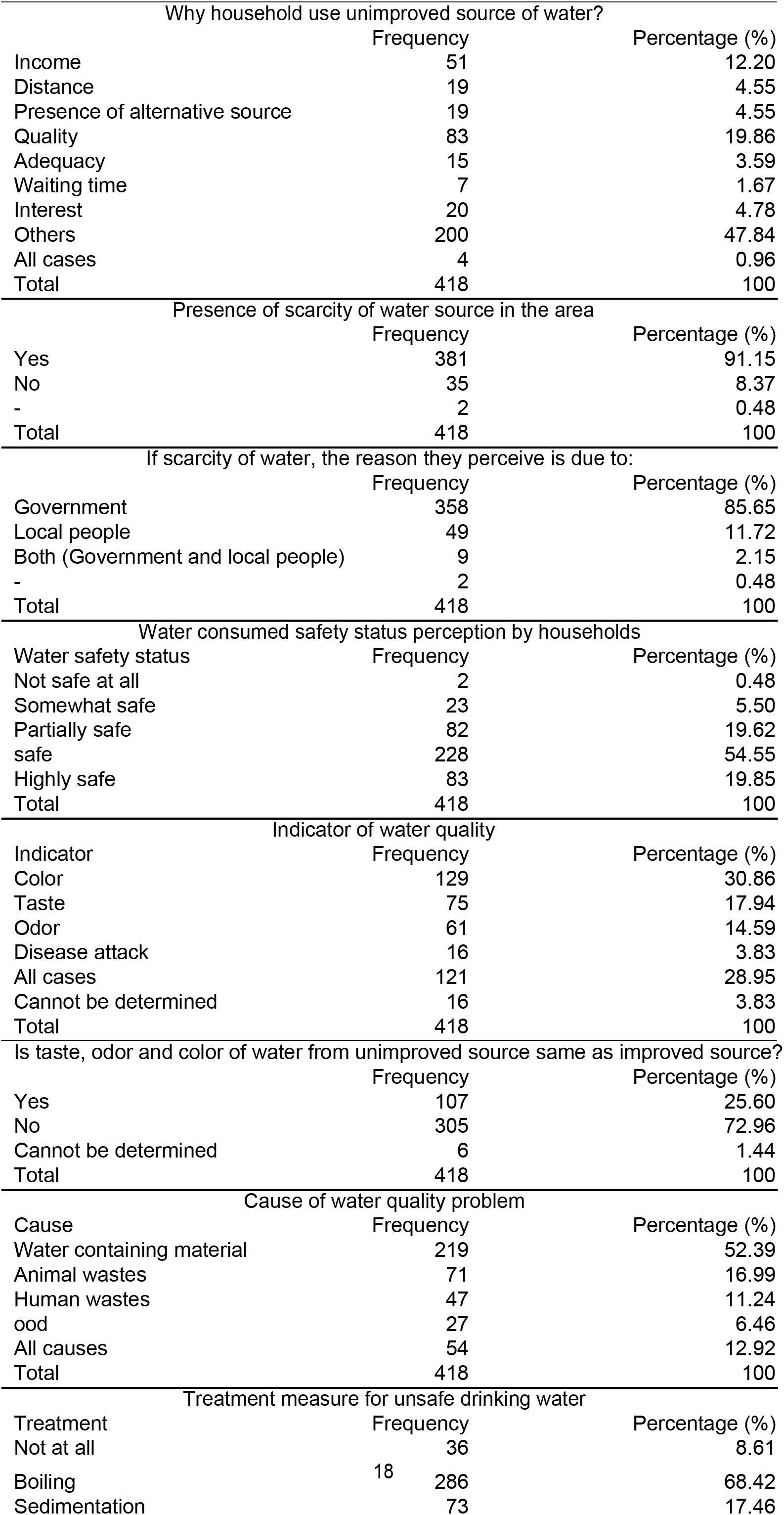
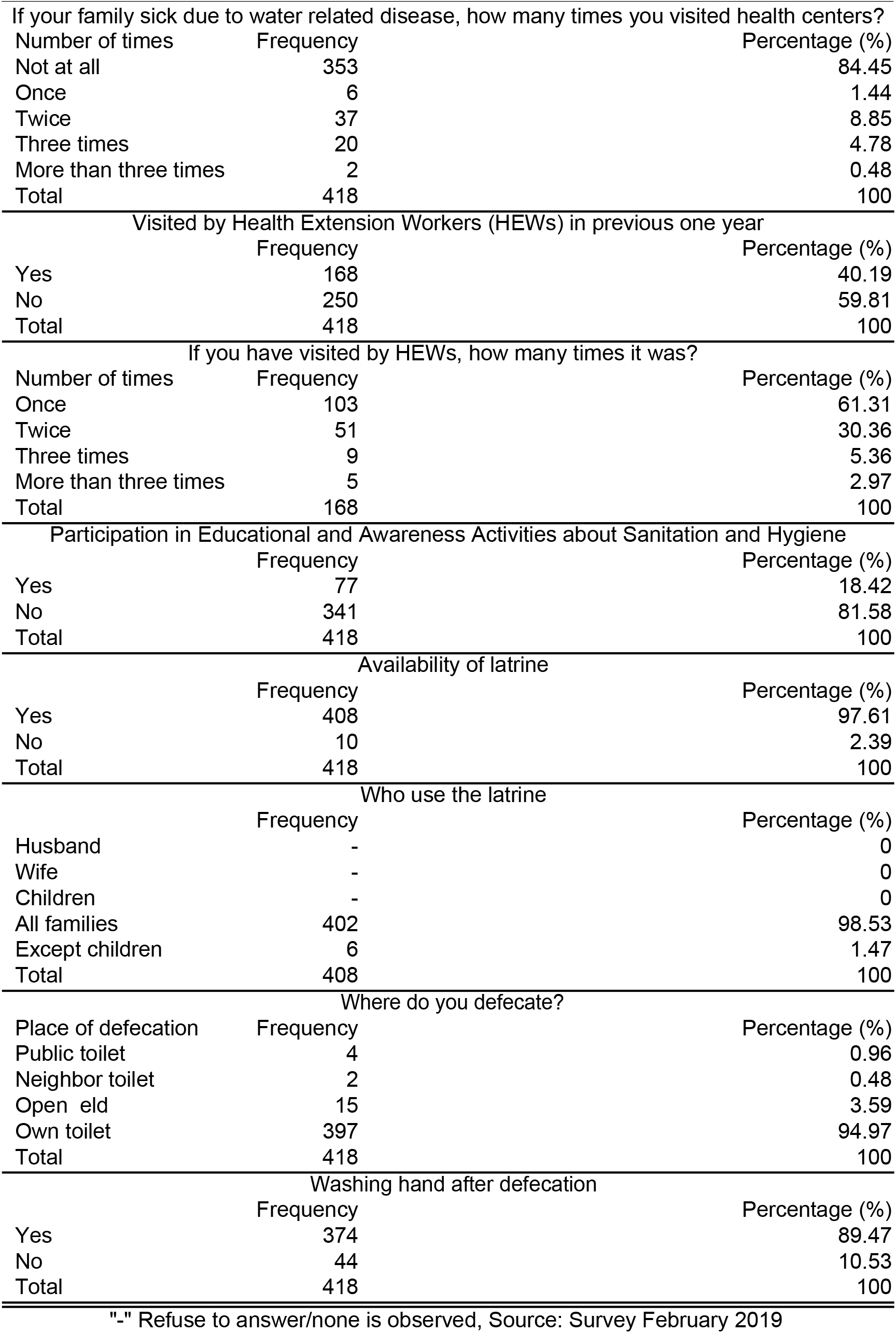
Summary Statistics of Water Source Using Practice, Quality and Sanitation Perception among Urban Households in North-West Ethiopia, 2019

**Table 2:**
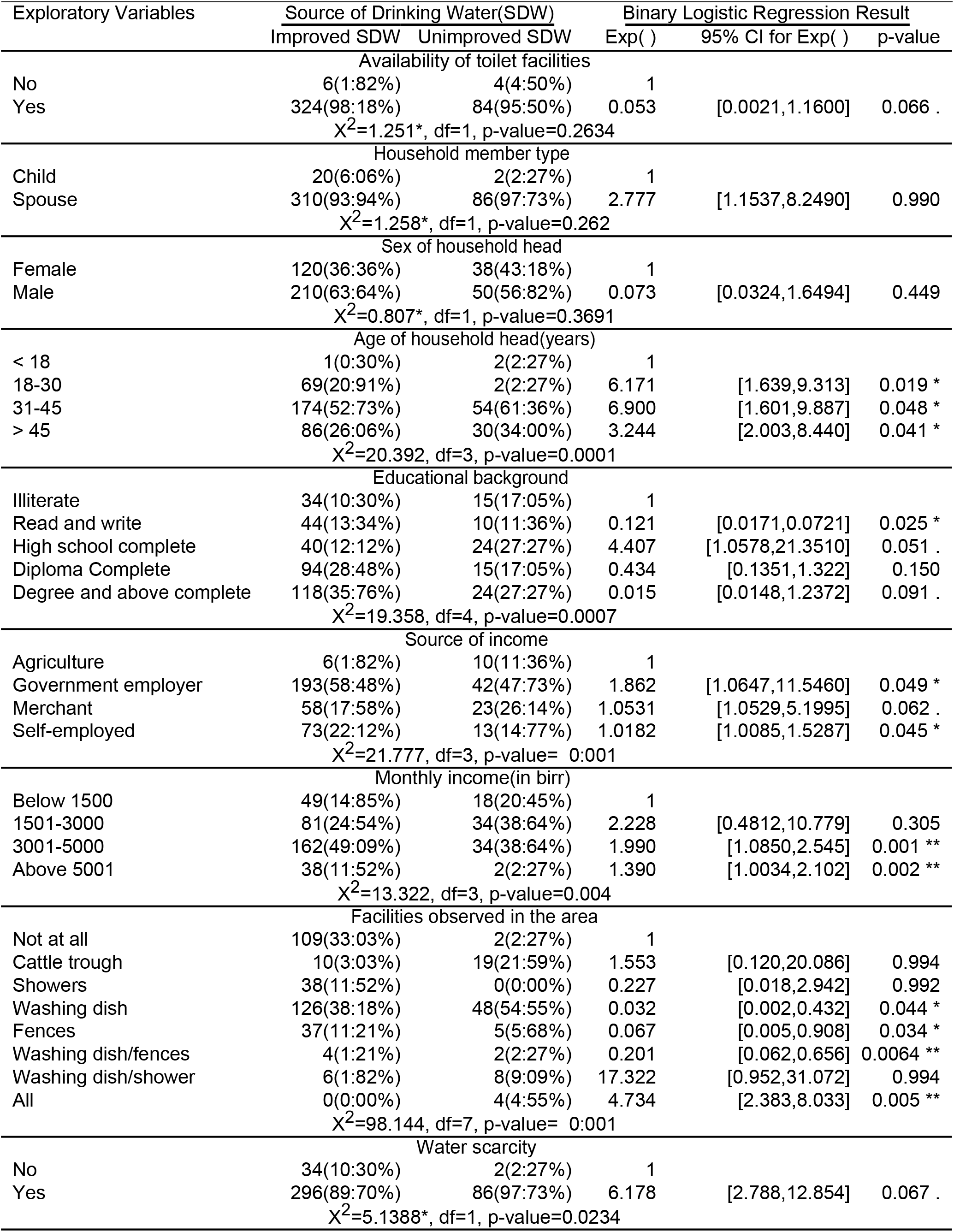
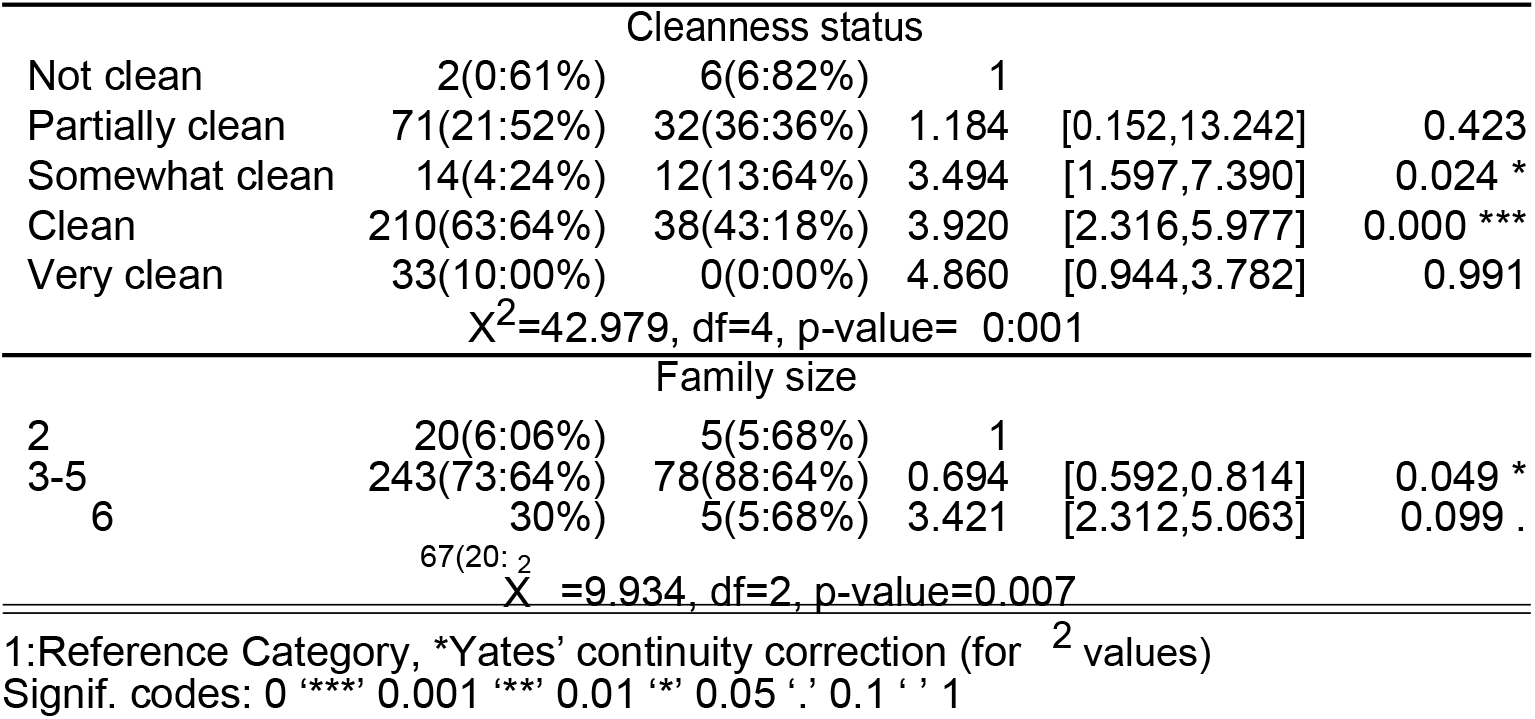
In uence of factors on SDW and Logistic regression of household SDW with the independent variables

### 3.1 Summary Statistics of Explanatory Variables

The summary statistics of the different explanatory variables used in this study are presented under (Table 1).

### 3.2 Unimproved Water source using Practice

Households use both improved and unimproved water sources for their daily water consumption. Based on the survey was undertaken in this study, about 330(78.95%) and 88(21.05%) households used improved and unimproved sources of water; respectively. Even, an improved source of water is in good quality it is not readily available. Sources of water (improved, unimproved) versus different explanatory variables were presented under (Table 2). (Table 1) shows the reason that household’s response to using unimproved source of water rather than an improved source of water. Households used unimproved source of water is due to income 51(12.20%), distance 19(4.55%), presence of alternative source 19(4.55%), quality 83(19.86%), adequacy 15(3.59%), waiting for time 7(1.67%), interest 20(4.78%), all cases 4(0.96%), and other (cases other than the listed) 200(47.84%) than the improved source of water The quality of the improved source of water is indeed better than the unimproved sources of water. About 83(19.86%) households preferred unimproved source of water than improved sources. It might be due to the accessibility of unimproved sources. Nearly half or 200(47.84%) households preferred unimproved sources than the cases of income, distance, presence of alternative source, quality, adequacy, waiting time, and interest. Other investigation to determine the factors (other than listed in this study) that households prefer unimproved sources than improved sources shall be undertaken.

Of 418 household respondents about 381(91.15%) perceive that water is scarce in the area; while 35(8.37%) respondents perceive as there is no scarcity of water. Two (0.48%) refuse to say about the presence or absence of the scarcity of water in the area. From this we can summarize that the scarcity of water is a serious issue, because above 90% households perceive that they live under scarce conditions. About 200(47.84%) households, the reason to use unimproved sources were “Others” (i.e., other than the listed). This might be due to the presence of scarcity of water from improved sources that households would like to use from unimproved sources.

If scarcity of water is available, the responded households asked about who would be responsible for. Of 418 households 358(85.65%) due to government, 49(11.72%) due to local people, 9(2.15%) due to both government and local people perceive that scarcity of water was occurred. About 2(0.48%) households who were refused to say about the presence or absence of the scarcity of water didn’t like to state the concerned body to be responsible for.

### 3.3 Water Safety Status and Quality Indicator Perception

The water safety status was presented by the Likert scale from low safety status to high safety status. The summary statistics (Table 1) shows the water consumed safety status of 418 responded households. About 83(19.85%) said that the water they consumed is highly safe, 228(54.55%) safe, 82(19.62%) partially safe, 23(5.50%) somewhat safe, and 2(0.48%) not safe at all; believed about the water they consumed. The majority of about 311(74.40%) households the water they consumed is safe and highly safe. Even if, the majority (three-fourth) of the respondents’ water safety status is safe, the remaining households should be targeted to improve their water safeness.

From the 418 responded households about 402(96.17%) had different perceptions about the quality of water being consumed from the different sources. About 129(30.86%) made complaints about the color, 75(17.94%) said it has taste, 16(3.83%) is attacked by disease, 61(14.59%) complained about its odor, 121(28.95%) all cases can be seen, while about 16(3.83%) can’t determine the water quality which they consume daily.

To know the general knowledge about the quality of water from improved and unimproved water sources, a question has been asked to households: “Is taste, odor, and color of the water from the unimproved source is the same as the improved source?” Of 418 respondents about 107(25.60%) answered “yes”, about 305(72.96%) answered “no”, while about 6(1.44%) answered as they cannot determine. The majority of 305(72.96%) households couldn’t differentiate the quality of water using taste, odor, and color either it is from improved or unimproved water sources. Even microorganisms are not ever identified by taste, odor, and color easily, using those identifiers to differentiate the water quality is cheap and fast in-door activity.

There are different causes of water quality problems from the source and in-door in accordance with the household reports. In-door, the water storage material takes the higher cause of the water quality problem. From the source animal and human wastes and floods are the main causes. (Table 1) provides the different causes of water quality problems of 418 households, about 219(52.39%) due to water containing material, about 71(16.99%) due to animal wastes, about 47(11.24%) due to human wastes, about 27(6.46%) due to flood, and about 54(12.92%) due to all cases.

Of 418 households about 286(68.42%) used boiling, about 73(17.46%) used sedimentation, about 2(0.49%) used chemical reagent, about 8(1.91%) used other treatment measures, about 13(3.11%) used all treatment measures (boiling, sedimentation, chemical); alternatively, while about 36(8.61%) didn’t use any treatment measures for unsafe drinking water. Majority 286(68.42%) households’ used boiling treatment measure is due to its undergone in-door, cheap, and easy. Modern treatment measures like using chemical reagents are not common and accessible for low-income households. There are individuals who don’t use any measure to treat unsafe drinking water.

From 418 households, about 353(84.45%) households’ families’ weren’t sick at all due to water-related disease in the previous 1 year. About 6(1.44%) household families were sick once due to water-related diseases and they had visited health centers; about 51(12.20%) household families were sick twice due to water-related diseases and of those about 37(8.85%) had visited health centers; about 6(1.44%) household families were sick three times and all had visited health centers, about 2(0.48%) household families were sick four and more time and all had visited health centers per year. Households, who were sick twice per year, had visited health centers three times. At the time, when the disease had gone to worse individuals visited health centers repeatedly. It is unusual for families who aren’t sick due to water-related diseases to visit health centers (i.e., all 353(84.45%) household families’, which weren’t sick due to water-related diseases hadn’t visited health centers for water-related disease).

(Table 1) shows the status of households, who were visited by health extension workers (HEWs) per year, of 418 households about 168(40.19%) had visited by HEWs, while about 250(59.81%) households hadn’t visited by HEWs in the previous year. Of about 168(40.19%) households, who had visited by HEWs, about 103(61.31%) visited once, about 51(30.36%) visited twice, about 9(5.36%) visited three times and about 5(2.97%) households had visited four and more times per year. More than half of the responded households hadn’t ever visited by HEWs. It shows poor management of the town health bureau. In the places where the communities live densely, communities need assistance for good health and quality of life to eradicate communicable diseases. Health extension workers (HEWs) play a great role in the development of community health and quality of life. Evidence shows HEWs have participated in the quality of life and family planning in the previous 15 years in the country.

### 3.4 Sanitation and Hygiene Practice

The participation of household members in educational and awareness activities about sanitation and hygiene had been played a great role in a healthy community and a clean environment. Of 418 households included in the survey about 77(18.42%) had been participated; while about 341(81.58%) hadn’t participated in educational and awareness activities concerning sanitation and hygiene in their locality. Community education and awareness activities about good health, quality of life (QoL), sanitation, and hygiene are given by health extension workers (HEWs) in the area. But, in the town only 168(40.19%) households had been visited by HEWs.

Regarding to the availability of latrine in the households, about 408(97.61%) households have latrine in compound and/or surrounding, for defecation; while about 10(2.39%) households didn’t have latrine. Of those who have latrine (408(97.61%) households), only about 402(98.53%) household toilets were accessible to all families. But about 6(1.47%) household toilets were accessible for families except children. About 397(94.97%) used own toilet, about 15(3.59%) used open-field, about 4(0.96%) used the public toilet and about 2(0.48%) used neighbors toilet for defecation. Open field toilet defecation was found within dense trees and drainage areas, which easily could disturb the surrounding environment and be eroded by the flood. Households use neighbors toilets are at a time when their home and/or safety tank are/is at the building stage(s).

The culture of household members washing their hand after defecation, of 418 households about 374(89.47%) households had washed their hand, while about 44(10.53%) households hadn’t adopted the culture of washing hands after defecation. Among 374(89.47%) households had washed their hand with water and additionally with soap, but inconsistently using soaps. The country also put a push e ort of washing hand after defecation by memorizing a day per year, nationally.

### 3.5 Predictors of Access to Improved Sources of Drinking Water

The chi-square test of association and the binary logistic regression results about the predictors of access to improved sources of drinking water were shown under (Table 2).

The results show that there was no significant association between the availability of toilet facilities and types of water sources used by residents of the town (x^2^ =1.251, df=1, p-value=0.2634). The availability of toilet facilities with improved sources was 98.18%(OR = 0.053; 95%CI : 0.0021-1.1600). There was no significant association of household member type (x^2^ =1.258, df=1, p-value=0.262) and the sex of the household head (x^2^ =0.807, df=1, p-value=0.3691) with the types of water sources used by residents of the town. Household member type being spouse was 93.94% (OR = 2.78; 95%CI: 1.1537-8.2490) to be accessible to improved sources of drinking water. The sex of the household head being male was 63.64% (OR = 0.073; 95%CI: 0.0324-1.6494) to be accessible to improved sources of drinking water. Based on the 95% CI for the odds ratio, of both household member type and sex of the household head, don’t significantly determine the source of water used by households.

There was a significant association between the age of the household head and the type of water source used by residents of the town (x^2^=20.392, df=3, p-value=0.0001). Age of household head (18-30 years) is 6.171 times higher than the age of being below 18 years; with 20:91% (95%CI: 1.639-9.313; p-value = 0.019). The accessibility of improved sources of drinking water are lower in the older age group (> 45 years) than medium age group (18-30 and 31-45 years), but better than the younger (below 18 years).

The educational status of household heads was significantly determined the presence of improved sources of drinking water within the households (x^2^=19.358, df=4, p-value=0.0007). Being able to read and write 0.121 times 13.34% (95%CI: 0.0171-0.0721), diploma complete 0.434 times 28.48% (95%CI: 0.1351-1.322); and degree and above complete 0.015 times 35.76% (95%CI: 0.0148-1.2372) were lower than illiterate household heads to be accessible to improved sources of drinking water. But, household heads with high school complete were 4.407 times 12.12% (95%CI: 1.0578-21.3510) higher than illiterate household heads.

There was a significant association between the main source of income of the households and the type of water sources used by the town residents (x^2^=21.777, df=3,p-value= 0:001). The main source of income of households’ through self-employer 22.12% (OR= 1.0182; 95%CI: 1.0082-1.5287), merchant 17.58% (OR= 1.0531; 95%CI: 1.0529-5.1995), and government employer 58.48% (OR= 1.862; 95%CI: 1.0647-11.5460) were significantly determine the source of water from improved sources with compared to households’ whose source of income were through agriculture. The accessibility of improved sources of drinking water were 86.2% increased for government-employed with compared to agriculture income households.

Additionally, there was a significant association between monthly income of households and the source of water used by the town residents (x^2^=13.322, df=3, p-value=0.004). Monthly income (1501-3000)(birr) was 24.54%(OR = 2.228; 95%CI : 0.4812-10.779); income (3001-5000)(birr) was 49.09%(OR = 1.990; 95%CI: 1.0850-2.545); and income above 5001 (birr) 11.52%(OR = 1.39; 95%CI: 1.0034-2.102) with compared to lower than 1500 (birr) income households. Facilities such as: - washing dish 38.18% (OR = 0.032; 95%CI: 0.002-0.432); fences 11.21% (OR = 0.067; 95%CI: 0.005-0.908), both washing dish and fences 1.21% (OR= 0.201; 95%CI: 0.062-0.656), and all facilities (OR = 4.734; 95%CI: 2.383-8.033) with compared to no facilities at all observed to determine the source of water from improved sources, but not cattle trough and showers; (x^2^=98.144, df=7,p-value=0.001).

There was a significant association between the cleanness status of the surrounding and the type of water used by the town residents (x^2^=42.979, df=4, p-value= 0.001). Somewhat clean 4.24% (OR=3.494, 95%CI: 1.597-7.390) and to be clean 63.64% (OR=3.92, 95%CI: 2.316-5.977) with compared to no clean surrounding were observed to determine the source of water from improved sources, but not partially clean and very clean surrounding cleanness status categories.

About Two-hundred ninety-six (89.70%) of households had a scarcity of water from improved sources. Nearly nine out of ten households were under a scarcity of water from improved sources. 6.178 times (odds) of water scarcity occurred from improved sources than water from non-improved sources (95%CI: 2.788 12:854) and (x^2^=5.1388, df=1,p-value=0.0234).

Households with large family size had accessible for water from improved sources. Family size (3-5) was 73.64% (OR = 0.694; 95%CI: 0.592-0.814), and family size (>=6) was 20.30% (OR = 3.421; 95%CI: 2.312-5.063) with compared to lower number of family size (<=2). From this we can conclude that medium number of family size (3-5) had got lower odds (decreased by 69.4%) to improve sources but a higher number of family size (>=6) had gained higher odds (3.42 times) to improved sources.

## 4 DISCUSSION

Different studies showed that many factors affect the supply of quality drinking water in households such as-age of household members [23–25], the gender of household members [23,26–34], occupation of the household head [35], improved and unimproved water sources in rural and urban areas [36–38,23,26], households standard of living (income) [39–45], education level of household members [33,46], household size and composition [26,47–50]. This study show the associated factors of households’ drinking water sources in urban households.

A study was undertaken in the area also concludes that nine of ten persons was under the problem of water scarcity; the supply was inadequate, and the quality was also low [67]. The current study finds out that about 78.95% population of the town were users of water from improved sources, while about 21.05% were used from unimproved sources. A study was undertaken in the surrounding rural areas of the Debre Tabor Town, Farta district shows about 57.10% of the population had access to improved water sources and the remaining from unimproved sources [66].

### Unimproved Water source using Practice

The population of the town used both improved and unimproved water sources for their daily consumption. Households use unimproved sources of water that were associated with several reasons such as-income, distance, presence of alternative sources, quality, adequacy, waiting for time, interest, and other cases. About 4.55% of the population due to distance and 19.86% of the population due to quality, used unimproved sources. The improved water sources in urban areas are located in short distances [51], and the quality of water is better from improved sources.

About 91.15% of the town’s population was under the problem of drinking water scarcity. It indicates that the supply is below 10%. The figure is lower with compared to 60% of the population has access to improved water sources in Africa and 42% water supply in sub-Saharan African countries [52]. About one billion population in the world has no access to safe and adequate water sources [53]; and the country report shows the presence of poor sanitation and drinking water infrastructure [13]. The lower supply of drinking water and the unimproved sanitation of households are associated [14]. It also affects the distribution of water in the area [15], and it leads to health risks [54–55]. EDHS report in 2016

[56] indicates that 97% of the urban population in Ethiopia had access to an improved source of drinking water, even if it was not clear about its quality [57]. But, in the current setting, the supply was below 10%. The report of [16] reason out that the problem is occurred due to drought and the Horn of Africa regional instability. The international report of [20] also suggested the push and pool factors of poor water and sanitation.

In the study area, 85.65% of the population perceived that the scarcity of water was associated with the local, regional and national government poor administration factors. This might coincide with the international report [20] of different factors, such as the absence of good drinking water infrastructure [13] and discontinuous supply of drinking water [15].

### Water Safety, Quality and Sanitation Perception

The presence of drinking water is vital for every human being. About 74.40% of the population in the study area consumed safe water, but about a quarter of the population consumed below a standard of safety. Even if, there is no pure water in nature [58], and about a billion people in the world don’t have access to safe drinking water [53], it is the right of citizens to get safe and accessible drinking water. Since unsafe water leads to water-borne diseases [59,40], the local government shall take this mandate to balance the right and responsibilities of its peoples. Those water-borne diseases are a major concern for households [35] and highly affect households from developing countries, who live in extreme conditions of poverty [61]. The risk of lack of safe water is more than any man-made destruction (such as-war, terrorism, and toxic weapons) [62]. Globally, millions of people die as a result of water-related diseases, WHO report [62].

Households use different perceptions to identify the quality of water they consume daily. They have used color, taste, disease attack, and odor. About 3.83% of the study population couldn’t determine the water quality they consumed. Households have got the water they consume from improved and unimproved sources. “Is the water perceptions are similar from those two sources?” was asked to households. About 25.60% of the population answered as similar perceptions were observed, but about 1.44% of the population answered as they couldn’t determine. About 72.96% of the population couldn’t differentiate the quality of water using taste, odor, and color either it was from improved or unimproved sources.

In the developing world, peoples don’t have access to safe water [53], and some defecate in an open-field [9]. Water sources contaminated with domestic and industrial wastes [13,62]. About half of the study population the causes for water quality was due to water containing material, poor indoor practices; 16.99% due to animal wastes and 11.24% due to human waste. In a place where water shortage is available, water may store for a long time. Water stored from 1 to 9 days increase the contamination by 67% [54]. In the current setting as investigators observed households stored water for more days. A study also shows that the town’s population had got water once per week [21]. The presence of animal and human wastes results in poor sanitation and hygiene. This leads to complicated water-related sickness and disease [55,40, 61–62].

When unsafe drinking water is observed, households use treatment measures such as-boiling, sedimentation and chemicals, and a combination of two or more. 68.42% of the study population were used boiling as a means of treatment measure and secondly, 17.46% were used sedimentation. This might be due to the lower cost. Unsafe drinking water is the cause of many water-borne diseases and leads to a health disorder.

About 84.45% of the study population hadn’t got sickness due to water-related diseases. This might be due to the cold weather condition of the area the distribution of the disease transmitter microorganisms or vectors is lower. The better experience observed in the area was, households had visited health centers during illness. Only 40.19% of the study population had visited by health extension workers (HEWs), which is lower than the country’s health extension coverage. It was not that much often HEWs had visited the households. It needs further investigation.

Communities participation in educational awareness activities in surrounding and/or the town sanitation and hygiene was/were very low, only 18.42%. Even if, the quality of latrine was not observed, but 97.61% of the population had been accessible for latrine facilities. The investigator recommends further study on the quality of latrine in the town.

A study undertaken in Ethiopia stated about 36% of the population practiced open-defecation [14], the current study shows 3.59%, which is lower. When the inaccessibility of water has occurred, people are forced to open-defecation [9]. Globally, open-defecation was declined from time to time [4]. In middle-income countries, 35% didn’t have water for handwashing with water and soap [20]. The current study shows only 10.53% is lower. Adequate water supply, good sanitation facilities, and proper hygiene practices improve the lives of the community [65]. The scarcity of water is associated with reduced sanitation facilities.

## 5 CONCLUSION

This study provides data on access, usage, and practices of water source among urban households in northwest Ethiopia. It provides a valuable insight into access to safe water and consequent demographic, socioeconomic conditions such as income, distances, and family size, sanitation and hygiene perceptions of households that can be associated with access to improved sources of water.

The major findings suggest that 78.95% of households used improved and 21.05% of households used unimproved water sources. Based on the report evidence, the study suggests as most developing countries, in Ethiopia, specifically in this study area, the scarcity of water, especially from improved source is very severe. The figure (91.15%) shows the town’s population is under the problem of water scarcity. Increasing demand with a population of safe and quality water puts in force the local governments to increase the supply. But, due to the lower supply of pure water to households, people put in force for using water from unimproved sources, which have a possibility to contaminate many infectious microorganisms and cause water-borne diseases. Even if, the presence of adequate drinking water is vital for humans, only 74.40% of the population consumes safe water, and the rest below the standard.

The causes of water quality for the population of 52.19% is due to the water-containing material, indoor practices, occurred under the availability of water shortages. Animal and human wastes are the second for the cause of water quality and can easily disturb the water sources. It is better to protect water sources from any contamination and use water treatment measures, when the water stored for a long time.

The study suggests the association of the types of sources of drinking water with the age of the household head, educational level of the household head, sources of income, monthly income, facilities observed, cleanness status of the surrounding, water scarcity and family size significantly determine the sources of water either it is from improved or unimproved sources, while; availability of toilet facility, household member type and sex of the household head are not significant. Thus, older headed households closely related to the availability of improved sources of drinking water. Educational status of the head of the household significantly determine the type of water source to be used. The type of source of income associate with the type of water source to be used in the households (i.e.,86.2%; 5.31% and 1.82%; for government employer, merchant and self-employed); respectively.

In the long-run Health Extension Workers (HEWs) shall be given attention for the improvement of the community sanitation and hygiene practices and give awareness. This might reduce the practice of open defecation.

In conclusion, the using of drinking water from improved sources was determined by different demographic, socio-economic, sanitation, and hygiene-related factors. Based on our investigation, being an older age group of the head of the household, being government employer, merchant and self-employed, being a higher income group, the presence of all facilities in the area, lived in a clean surrounding and lower family size are the determinant factors of using drinking water from improved sources. It is recommended that the local, regional, and national governments and other supporting organizations shall improve the accessibility and adequacy of drinking water from improved sources through short and long time plan for the well-being of the community in the area.

## 6 Limitations

This study was conducted with the data collected from the households’ perceptions about the source of water use, water quality, and sanitation and hygiene practices in the area. Thus, the current study didn’t undertake a verifiable water quality measures.

## Abbreviations

CI: Confidence Interval
CSA: Central Statistics Agency
DTT: Debre Tabor Town
EDHS: Ethiopian Demographic and Health Survey
HEW: Health Extension Worker
ICF: International Children’s Fund
JMP: Joint Monitoring Program
l/p/d: liters per person per day
OR: Odds Ratio
SDW: Sources of Drinking Water
SNNP: Southern Nations and Nationalities of Peoples
UNICEF: United Nation International Children Fund
WHO: World Health Organization

## Declarations

### Ethics approval and consent to participate

This study was reviewed and approved by Debre Tabor University Research Ethics Committee and data collection were possible after written permission were found from study participants.

### Consent to publish

Not applicable.

### Availability of data and materials

Not applicable. All the data supporting the findings are contained within the manuscript.

### Authors’ Contributions

SG conducted a literature search, planned the study, carried out data collection, performed data analysis and interpretation and drafted the manuscript. Authors EY, BD and AD reviewed the literature search, design of the study, data analysis and data interpretation. All authors read and approved the final manuscript.

## Acknowledgments

The authors wish to thank Debre Tabor University, study participants, and data collectors for making this investigation is possible.

